# Impact of repeated short light exposures on sustained pupil responses in an fMRI environment

**DOI:** 10.1101/2023.04.12.536600

**Authors:** Elise Beckers, Islay Campbell, Roya Sharifpour, Ilenia Paparella, Alexandre Berger, Jose Fermin Balda Aizpurua, Ekaterina Koshmanova, Nasrin Mortazavi, Puneet Talwar, Siya Sherif, Heidi I.L. Jacobs, Gilles Vandewalle

## Abstract

Light triggers numerous non-image forming (NIF), or non-visual, biological effects. The brain correlates of these NIF effects have been investigated, notably using Magnetic Resonance Imaging (MRI) and short light exposures varying in irradiance and spectral quality. However, it is not clear whether having light in subsequent blocks may induce carry over effects of one light block onto the next, thus biasing the study. We reasoned that pupil light reflex (PLR) was an easy readout of one of the NIF effects of light that could be used to address this issue. We characterized the sustained PLR in 13 to 16 healthy young individuals under short light exposures during three distinct cognitive processes (executive, emotional and attentional). Light conditions pseudo-randomly alternated between monochromatic orange light [0.16 melanopic Equivalent Daylight Illuminance (mel EDI) lux] and polychromatic blue-enriched white light of three different levels [37, 92, 190 mel EDI lux]. As expected, higher melanopic irradiance was associated with larger sustained PLR in each cognitive domain. This result was stable over the light block sequence under higher melanopic irradiance levels as compared to lower ones. Exploratory frequency-domain analyses further revealed that PLR was more variable within a light block under lower melanopic irradiance levels. Importantly, PLR varied across tasks independently of the light condition pointing to a potential impact of the light history and/or cognitive context on PLR. Together, our results emphasize that the distinct contribution and adaptation of the different retinal photoreceptors influence the NIF effects of light and therefore potentially their brain correlates.

## Introduction

Besides its visual function, light affects numerous so-called non-imaging-forming (NIF) or non-visual biological processes, such as the entrainment of circadian rhythms, the regulation of body temperature, the constriction of the pupil, the secretion of hormones, and the stimulation of alertness and cognition (Cajochen et al., 2005; Fisk et al., 2018; Gamlin et al., 2007; Lok et al., 2018). The intrinsically photosensitive retinal ganglion cells (ipRGCs), expressing the photopigment melanopsin, constitute a distinct class of retinal photoreceptors heavily involved in mediating the NIF impacts of light (Berson et al., 2002; Lucas et al., 2014). Depending on the light level, ipRGCs also receive inputs from rods and cones that are added to their intrinsic response to modulate the activity of their brain projections (Güler et al., 2008). The sensitivity of melanopsin is maximal for blue wavelength light, at about 480nm, such that the overall sensitivity of ipRGCs and NIF responses is shifted towards the shorter wavelength portion of the visible light spectrum, around 460-480nm (Brainard et al., 2001; Thapan et al., 2001).

The diversity of the NIF impacts of light is reflected in the widespread and complex projections of ipRGCs to numerous subcortical regions, including the suprachiasmatic nucleus (SCN), site of the master circadian clock, the ventrolateral preoptic nucleus (VLPO) involved in sleep regulation, and the olivary pretectal nucleus (OPN) playing a key role in pupil light reflex (PLR) (Hattar et al., 2006). These projections were mainly identified in rodents and translation to humans is not straightforward, making the exact neural mechanisms underlying the NIF impacts of light still insufficiently understood in humans. Over the past two decades, non-invasive techniques such as functional magnetic resonance imaging (fMRI) have started elucidating part of the brain mechanisms underlying the stimulating effect of blue wavelength light on human cognition. These studies led to the idea that this activating effect was mediated through subcortical areas involved in alertness and sleep regulation, and cortical brain regions, in a time- and task-dependent manner (Gaggioni et al., 2014; Vandewalle et al., 2009). Most of these fMRI studies used repeated short light exposures (1 minute or less) alternating between different spectral compositions [e.g. (Daneault et al., 2014; McGlashan et al., 2021; Vandewalle et al., 2011; Vandewalle et al., 2007; Vandewalle et al., 2010)]. Whether the effect of light in one block carries over the next one, thus potentially biasing these fMRI results, remains unclear (e.g., exposure to 40s of monochromatic blue light following 20s of darkness following 40s of monochromatic green light).

Here, we addressed this issue by simultaneously acquiring fMRI and pupillometry data. PLR consists of the constriction of the pupil in response to an increased illumination of the retina and constitutes therefore an easy readout of a NIF impact of light. PLR is driven by the combined contribution of rods, cones, and ipRGCs intrinsic response (Gamlin et al., 2007). Rods and cones play a primary role at low irradiance levels and/or over the earliest part of the illumination, while the intrinsic response of ipRGCs has a dominant contribution under relatively higher irradiance levels and/or following the initial period of illumination. While pupil constriction is robustly affected by light, it shows signs of progressive adaptation with a gradual pupil dilation in continuous exposure with a rate of change dependent on the irradiance level (Gooley et al., 2012).

Importantly, the diameter of the pupil can also fluctuate over prolonged periods independently from changes in luminance through ongoing cognitive activity (Joshi & Gold, 2020). However, whether these prolonged fluctuations impact PLR as a function of the cognitive context is not known. Interestingly, these non-luminance fluctuations may be driven in part by the tonic activity of the locus coeruleus (LC)-noradrenergic system, which is suggested as an important region of the brainstem mediating part of the NIF impacts of light (Joshi et al., 2016; Megemont et al., 2022). Quantification of these fluctuations under various light exposures may therefore inform on the activity of the LC underlying the NIF responses to light.

In the present study, we sought to characterize the sustained pupil response during an fMRI protocol including different cognitive tasks under short light exposures. Participants were alternatively exposed to short blocks of different irradiance levels, expressed in melanopic (mel) Equivalent Daytime Illuminance (EDI) lux: in darkness (<0.1 lux), exposed to a low-level monochromatic orange light (0.16 mel EDI lux), or 3 intensities of blue-enriched polychromatic white light (6500K; 37, 92, and 190 mel EDI lux). While under light, participants completed executive, emotional, and attentional tasks, respectively lasting 25, 20 and 15 minutes. We aimed to replicate the larger sustained PLR under higher irradiance levels and assess whether PLR was stable across tasks and time. Under higher melanopic irradiance levels, we expected a stronger PLR not modulated by cognitive processes or protocol time. We also conducted exploratory frequency-domain analyses on PLR variability under different light conditions and cognitive contexts to potentially relate them to LC activity (Nguyen et al., 2022).

## Methods

### Participants

In total, twenty-two healthy young adults (15 females; age 23.3y ± 4.3) were recruited to take part in this study after providing written informed consent. All participants were screened via semi-structured interviews and clinical questionnaires to assess exclusion criteria such as history of major psychiatric or neurological disorders, sleep disturbances and extreme chronotypes. They scored within normal ranges on the 21-item Beck Anxiety Inventory (Beck et al., 1988), the Beck Depression Inventory II (Beck et al., 1961), the Epworth Sleepiness Scale (Johns, 1991), the Horne-Östberg questionnaire (Horne & Ostberg, 1976), the Pittsburgh Sleep Quality Index (Buysse et al., 1989), and the Seasonal Pattern Assessment Questionnaire (Rosenthal, 1984).

All participants were non-smokers, eligible for MRI scanning, and free of psychoactive medications. Although no thorough ophthalmologic examination was performed, none of the participants reported ophthalmic disorders or color blindness. All participants reported normal hearing abilities. We excluded participants with a Body Mass Index (BMI) above 28, excessive caffeine (>4 caffeine units/day) or alcohol consumption (>14 alcohol units/week), travelling through more than one time zone during the last two months or working on non-regular office hours. Women were not pregnant or breastfeeding. Due to the exclusion of data sets with bad or missing data, the analyses of the executive, emotional and attentional tasks included 16, 13 and 16 participants, respectively. A summary of participants’ characteristics respective to each task can be found in **Table 1**.

**Table 1.** Study sample characteristics. Characteristics of the total study sample, and of the participants included in the executive, emotional, and attentional tasks, respectively. BMI: Body Mass Index. The education level is computed as the number of successful years of study. Scores of the BAI (Beck Anxiety Inventory), BDI-II (Beck Depression Inventory II), ESS (Epworth Sleepiness Scale), HO (Horne-Ostberg), PSQI (Pittsburgh Sleep Quality Index) and SPAQ (Seasonal Pattern Assessment Questionnaire). Average value ± standard deviation (SD). F: Female, M: Male.

|  | ***Total sample***<br>***(n = 22)*** | ***Executive task***<br>***(n = 16)*** | ***Emotional task***<br>***(n = 13)*** | ***Attentional task***<br>***(n = 16)*** |
| --- | --- | --- | --- | --- |
| *Age (years)* | 23.27 ± 4.33 | 23.25 ± 4.7 | 23.92 ± 4.66 | 24.13 ± 4.1 |
| *BMI (kg/m<sup>2</sup>)* | 21.38 ± 2.52 | 20.93 ± 2.34 | 21.48 ± 1.65 | 21.68 ± 2.31 |
| *Education (years)* | 14.2 ± 3.14 | 14.2 ± 3.45 | 14.45 ± 2.58 | 15.07 ± 2.43 |
| *BAI* | 6.25 ± 5.79 | 5.93 ± 6.17 | 8.36 ± 6.58 | 6.33 ± 4.59 |
| *BDI-II* | 6.7 ± 5.53 | 5.53 ± 5.8 | 7.45 ± 5.89 | 6.73 ± 5.42 |
| *ESS* | 6.4 ± 3.1 | 5.93 ± 3.06 | 7.09 ± 3.53 | 6.33 ± 2.69 |
| *HO* | 47.45 ± 9.34 | 48.07 ± 8.06 | 46.64 ± 6.47 | 48.13 ± 9.76 |
| *PSQI* | 4.4 ± 2.76 | 4.27 ± 2.89 | 4.73 ± 3.29 | 4.4 ± 3.02 |
| *SPAQ* | 0.95 ± 0.83 | 1 ± 0.85 | 1.09 ± 0.83 | 1 ± 1.07 |
| *Sex* | 15 F – 7 M | 11 F – 5 M | 9 F – 4 M | 11 F – 5 M |

The protocol was approved by the Ethics Committee of the Faculty of Medicine of the University of Liège and all participants received monetary compensation for their participation.

### Protocol

All participants followed a loose sleep-wake schedule for a 7 day period preceding the experiment at their habitual sleep and wake-up time (+/-1 hour) to avoid excessive sleep restriction while maintaining uniform realistic life conditions. Adhesion to the pre-defined schedule was verified through wrist actimetry (AX3 accelerometer, Axivity, United Kingdom) and sleep diaries. Participants were asked not to take nap during this period. Volunteers were requested to refrain from all caffeine and alcohol-containing beverages, and extreme physical activity for at least 3 days before participating in the study. The experiment took place either in the morning (N = 18) or in the evening (N = 4) intending to investigate the time-of-day effect of light exposure on brain functions and behavior in the future. Although controlled for in statistical analyses, the later time-of-day aspect will not be considered in the present paper. Data acquisitions took place between December 2020 and May 2022. On the day of the experiment, participants arrived in the laboratory 1.5 to 2 hours after their habitual wake-up time or 1.5 to 2 hours before their habitual bedtime and were exposed to 5 minutes of bright white light (∼1000 lux) followed by 45 minutes of dim light (<10 lux) to control for recent light history. During the dim light adaptation period, detailed instructions were given regarding the study, MRI environment and cognitive tasks to be performed in the MR scanner (**Fig. 1**). Task practices were also completed on a laptop aiming for an accuracy score of at least 75%.

**Figure 1.**
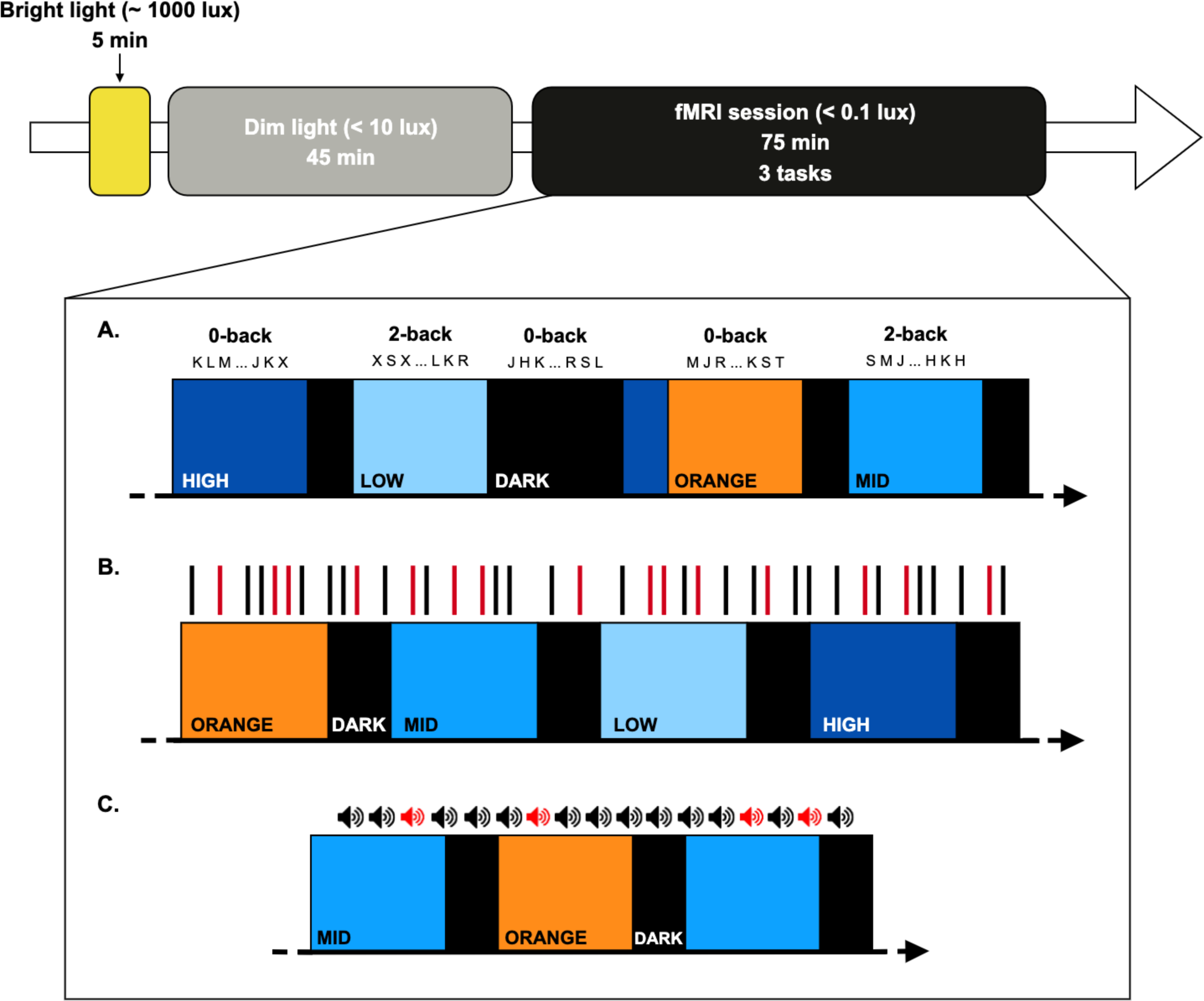
Graphical representation of the experimental protocol. Following a 7 day period of loose sleep-wake schedule (verified through wrist actimetry and sleep diaries), participants arrived at the laboratory 1.5 to 2 hours after their wake-up time, or 1.5 to 2 hours before their bedtime. After a light adaptation period (5 min of bright light (∼1000 lux) followed by 45 min of dim light (< 10 lux)), participants completed 3 auditory cognitive tasks during a functional magnetic resonance imaging (fMRI) session. Tasks respectively probed executive, emotional and attentional processes. While the protocol always started with the executive task, the order of the emotional and attentional tasks was counter-balanced across participants. **(A)** Detailed experimental design for the executive task. The task consisted of a two-level variant of the N-back task: 0-back and 2-back. Participants had to detect whether the current item matched the predefined letter “K” (0-back), or whether the current item was identical to the letter presented 2 items earlier (2-back). Tasks blocks included 15 items and lasted 30s. While performing the task, participants were exposed to pseudo-randomly alternating polychromatic white light of three different intensities (LOW: 37, MID: 92, HIGH: 190 mel EDI lux; 6500K) and a monochromatic orange light (0.16 mel EDI lux; 589nm). In total, 11 blocks for each light condition were included. Each light block lasted 30-67s and were interleaved by short 10s rest periods. **(B)** Detailed experimental design for the emotional task. The task consisted of a pure gender discrimination of auditory vocalizations while being exposed to the pseudo-randomly alternating polychromatic white light of three different intensities (LOW: 37, MID: 92, HIGH: 190 mel EDI lux; 6500 K) and a monochromatic orange (0.16 mel EDI lux; 589nm) light. 5 blocks of each light condition were included. Each light block lasted 30-40s and was followed by 20s period of darkness. Untold to the participants, vocalizations were pronounced with angry (red bars) and neutral (black bars) prosody, pseudo-randomly and equally distributed across the four light conditions. **(C)** Detailed experimental design for the attentional task. The task consisted of the detection of rare deviant tones (20%), within a stream of frequent standard tones (80%). Whilst completing the task participants were pseudo-randomly exposed to a polychromatic white light (MID: 92 mel EDI lux; 6500K) and a monochromatic orange light (0.16 mel EDI lux; 589nm). 7 blocks for both light conditions were included. Each block lasted 30s and was followed by a 10s period of darkness. Standard (black) and deviant (red) stimuli were equally distributed across the two light conditions.

### Light exposure

In the MRI, light was administered through a computer-controlled MR-compatible set-up designed in-lab and consisting of 3 main parts. First, a polychromatic blue-enriched white LED light source (SugarCUBE, Ushio America, CA, USA) with various intensities; second, a motor-driven filter wheel (AB300-Series, Spectral Products, NM, USA) allowing the automated changes in light conditions respectively using an Ultra Violet (UV) long bypass filter (433-1650nm) or a monochromatic orange light filter (589nm); third, an 8-meter-long metal-free dual-end optic fiber (Setra Systems, MA, USA) transmitting the light to participants’ eyes. A stand placed at the back of the head coil allowed the reproducible fixation and orientation of the optic fiber ends towards the inside of the coil and created a relatively uniform and indirect illumination towards participants’ eyes.

While performing functional tasks in the MR environment, participants were either maintained in darkness (<0.1 lux) or exposed to short light blocks which could be of 4 types, varying in irradiance level and spectral composition. Light pseudo-randomly alternated between a monochromatic orange light (4.24×10^12^ photons/cm^2^/s; 589nm, 10nm at full-width half maximum; 0.16 mel EDI lux) and a polychromatic LED light enriched in blue wavelengths of three different irradiance levels (6500K; 37, 92 and 190 mel EDI lux). Light spectra and light characteristics can be found in **Figure 2** and **Table 2**, respectively.

**Figure 2.**
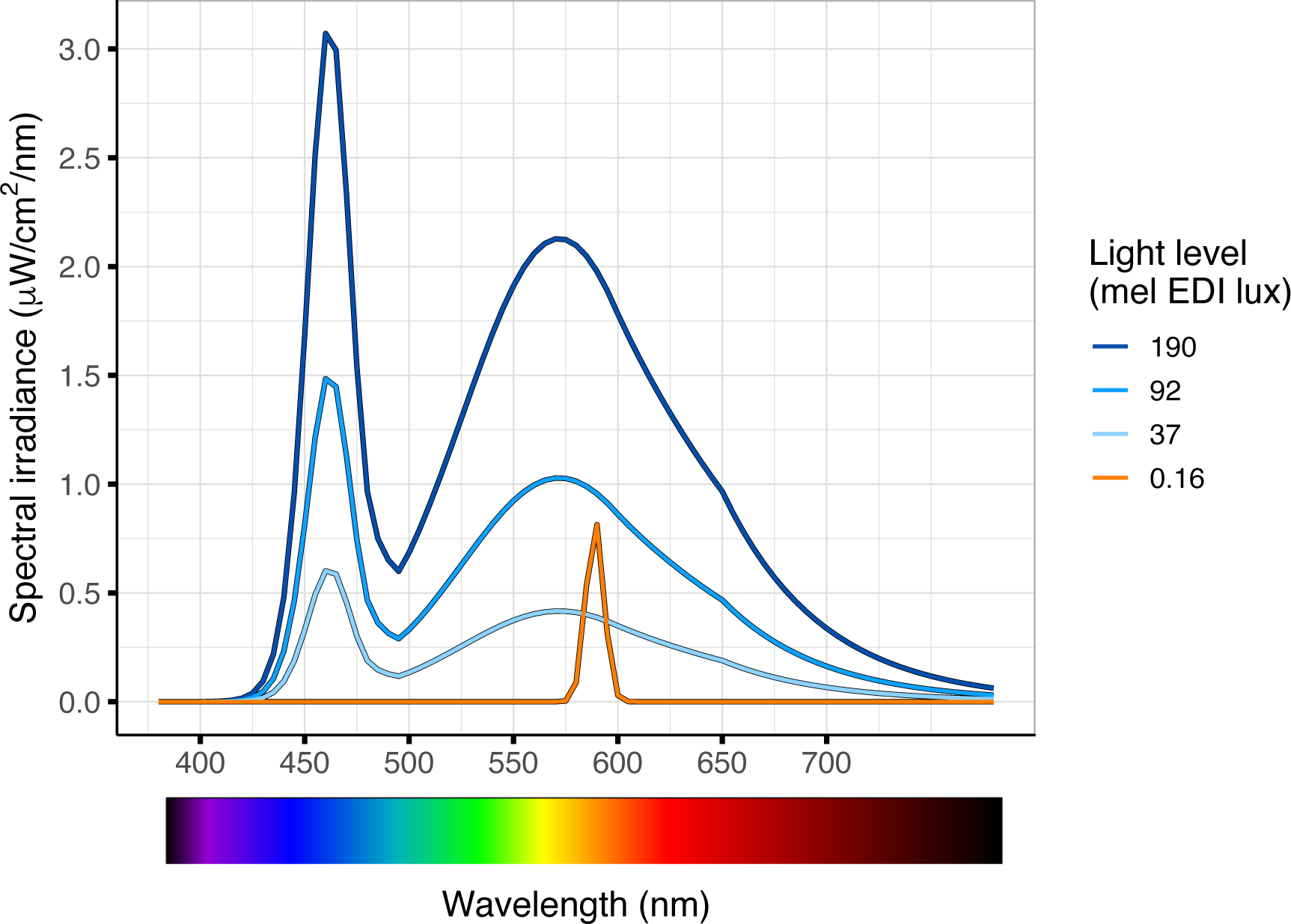
Spectrum power distribution of the four different light conditions. Monochromatic orange light (0.16 mel EDI lux), 589nm; Polychromatic white LED light enriched in blue wavelengths of three distinct irradiance levels (37, 92, 190 mel EDI lux; 6500K). *Adapted from Campbell et al. (bioRxiv)*

**Table 2.** Light characteristics of the four light conditions. Characteristics of polychromatic blue-enriched white light of 3 irradiance levels (LOW: 37 mel EDI lux, MID: 92 mel EDI lux, HIGH: 190 mel EDI lux) and monochromatic orange light (589nm).

|  | **LOW** | **MID** | **HIGH** | **Orange** |
| --- | --- | --- | --- | --- |
| **Lux** | 47 | 116 | 240 | 7.5 |
| **Peak Spectral Irradiance (nm)** | 460 | 460 | 460 | 590 |
| **Melanopic EDI (ipRGCs)** | 37 | 92 | 190 | 0.16 |
| **Rhodopic EDI (Rods)** | 39 | 97 | 201 | 0.94 |
| **Cyanopic EDI (S-cones)** | 32 | 79 | 163 | 0 |
| **Chloropic EDI (M-cones)** | 44 | 110 | 227 | 5 |
| **Erythropic EDI (L-cones)** | 46 | 113 | 233 | 8 |
| **Irradiance ($\mu\text{W}/\text{cm}^2$)** | 15 | 36 | 74 | 1.4 |
| **Photon flux($1/\text{cm}^2/\text{s}$)** | 4.12E+13 | 1.02E+14 | 2.10E+14 | 4.24E+12 |
| **Log Photon Flux ($\log_{10}$)**<br>**($1/\text{cm}^2/\text{s}$)** | 13.61 | 14.01 | 14.32 | 12.63 |
| **Narrowband peak** | - | - | - | 589 |
| **Narrowband FWHM** | - | - | - | 10 |

### Auditory cognitive tasks

The fMRI session included 3 auditory tasks probing cognitive functions such as executive, emotional, and attentional processes lasting about 25, 20, and 15 minutes, respectively. **Figure 1** depicts an overview of the global protocol. The executive task consisted of an auditory variant version of the n-back task (Collette et al., 2005) with two levels presented in distinct blocks: 0-back and 2-back. Participants had to detect whether the current item matched the predefined letter “K” (0-back), or whether the current item was identical to the letter presented 2 items earlier (2-back). Each task block lasted 30s and therefore included 15 items. Blocks and task levels were pseudo-randomly presented across all light conditions. Overall, the executive task included 11 blocks of each of the 4 melanopic irradiance levels. Light blocks lasted between 30 and 67s, and were interleaved with short 10s rest periods.

During the emotional task, participants were asked to indicate the gender of meaningless auditory vocalizations, while ignoring the negative and neutral prosodies of these stimuli (Banse & Scherer, 1996). In total, 240 auditory stimuli were pronounced by professional actors (50% female). Tasks events were pseudo-randomly and equally spread over the four light conditions. The emotional task included 5 blocks for each of the four melanopic irradiance levels. Blocks of light lasted 30 to 40s and were interleaved by 20s of darkness.

The attentional task was a mismatch negativity, or oddball task (Stevens et al., 2000). Participants had to report the detection of rare deviant tones (20%, 1000Hz, 100ms) within a stream of frequent standard tones (80%, 500Hz, 100ms). To maintain task duration acceptable for the participants and below 15 min, only two light conditions were included in the attentional task: one level of polychromatic, blue-enriched LED light (6500K; 92 mel EDI lux), and the monochromatic orange light. In total, 315 tasks events were pseudo-randomly spread over the two melanopic irradiance levels. The attentional task included 7 blocks for both light conditions. Participants were exposed to 30s of light blocks interleaved by 10s of darkness.

While the protocol always started with the executive task, the order of the emotional and attentional tasks was pseudo-randomized across participants. Auditory stimuli and instructions were delivered through MR-compatible earplugs (Sensimetrics, Malden, MA) controlled via a computer running OpenSesame software (version 3.2.8) (Mathôt et al., 2012). Prior to the start of the experiment, a volume check was performed to ensure the scanner noise was not undermining a proper perception of auditory stimuli. Responses to fMRI tasks were collected through an MR-compatible button box (Current Design, Philadelphia, PA) placed in the participants’ dominant hand. At the end of each task, participants stayed in near darkness for about 5 min which were used to acquire a control MR sequence, recalibrate the eye tracking system, and repeat instructions to the participant.

### Data acquisition

Data were acquired while participants were lying in a 7T MAGNETOM Terra MR scanner (Siemens Healthineers, Erlangen, Germany) with a 32-channel receiver and 1-channel transmit head coil (Nova Medical, Wilmington, MA, USA). Pupil size was continuously measured using an MR-compatible infrared eye tracking system at a sampling rate of 1000 Hz (EyeLink 1000Plus, SR Research, Ottawa, Canada) with a monocular recording (right pupil was used). The eye tracking system returned pupil area as an arbitrary unit, i.e., the number of pixels considered part of the detected pupil. Before the execution of each task, a pupil calibration was performed. Besides acquiring pupil data, the eye tracking system enabled the constant monitoring of participants’ sleepiness.

### Data analysis

### Pupil signal preprocessing

Pupil data analyses were conducted offline in MATLAB R2019b (MathWorks, MA, USA) where the data was cleaned at first. Identified blinks were replaced using linear interpolation and data were smoothed using the *rlowess* built-in robust linear regression function. Data sets with more than 25% of missing or corrupted data were excluded from the analysis.

Since we were interested in sustained PLR, the first 2s of each light block were discarded prior to averaging pupil value per light block. Pupil size averages were normalized with respect to the average pupil size during the darkness periods prior to averaging per light condition. For frequency domain analyses, power spectral density (PSD) was estimated via Welch’s method through the built-in *pwelch* function on 4-second rectangular windows with a 50% overlap, excluding missing values. In line with previous studies, the frequency band of interest was set between 0.5 and 4Hz with 0.5Hz sensitivity (Joshi et al., 2016; Nakayama & Shimizu, 2021; Peysakhovich et al., 2015). Total power per light condition between 0.5 and 4Hz was first computed by summing PSD values in this frequency range, prior to being normalized to the total power under darkness periods, and then converted to dB through a log_10_ scaling.

### Statistics

Generalized linear mixed models (GLMM) implemented in SAS 9.4 (SAS Institute Inc, NC, USA) were used with averaged normalized pupil value as the dependent variable, subject as random effect, and melanopic irradiance, task, and block order (when applicable) as repeated measures [autoregressive (1) correlation], while adjusting for sex, age, BMI, and time of day. The statistical significance threshold was set at *p*<.05. The distribution of the dependent variable was assessed prior to each model and the GLMM was set accordingly. Cook’s distance > 1 was used as cut-off to detect outlier values. No outliers were detected in the analyses of the executive, emotional and attentional tasks. Partial R^2^ (R^2^*) values were computed to estimate the effect sizes of significant effects in each model (Jaeger et al., 2017). The first implemented model tested for simple effects, along with interaction effects, of melanopic irradiance and task nature on the total averaged normalized pupil size while considering the three tasks altogether. Post-hoc analyses were conducted on the task nature while using a Tukey adjustment. Then, a second model tested for simple effects of melanopic irradiance in each task separately and post-hoc analyses were conducted on melanopic irradiance levels. In order to investigate the pupil response stability across light blocks for each task, the averaged normalized pupil size from the first and the last block of each light condition were tested for simple effects and interaction against melanopic irradiance and block order. Post-hoc analyses were conducted on this interaction and results were corrected for multiple comparisons using a Tukey adjustment.

In the scope of exploratory frequency-domain analyses, a fourth GLMM was implemented with total normalized PSD as the dependent variable, subject as random effect, and melanopic irradiance as a repeated measure [autoregressive (1) correlation], while adjusting for sex, age, BMI, and time of day. For the executive task, one dataset was reported as an outlier. As a consequence, the analysis of the executive, emotional and attentional tasks included 15, 13 and 16 participants, respectively. Results from models including and excluding the outlier were not differing. This model tested for simple effects of melanopic irradiance on the total normalized PSD in the 0.5-4Hz frequency range. Post-hoc analyses were conducted on melanopic irradiance levels and results were corrected for multiple comparisons using a Tukey adjustment.

## Results

### Melanopic EDI lux level-dependent sustained pupil response

Sustained averaged pupil response was first related to melanopic irradiance levels (0.16, 37, 92 and 190 mel EDI lux) considering all cognitive contexts together. The GLMM yielded a significant main effect of the irradiance level (F_(3,37)_=530.6, *p*<.0001, R^2*^=.98), such that higher melanopic irradiance level was associated with smaller pupil size. Importantly, a significant main effect of the task (F_(2,52)_=3.8, *p*=.03, R^2*^=.13) was detected, but, critically, no significant interaction effect between task and melanopic irradiance (F_(4,37)_=0.85, *p*=.5). Post-hoc analyses revealed no significant difference in pupil response between the emotional and the attentional tasks (t_55_=-0.09, *p*=.996), while statistical trends were detected between the executive task and both the emotional (t_63_=2.38, *p*=.053) and attentional tasks (t_55_=2.1, *p*=.099) such that the PLR was suggested to be reduced in the executive task as compared to the two other tasks.

Then, taking each task individually, a significant main effect of the melanopic irradiance level was also observed (Executive: F_(3,15)_=289.8, *p*<.0001, R^2*^=.98; Emotional: F_(3,12)_=165.1, *p*<.0001, R^2*^=.98; Attentional: F_(1,6)_=332.5, *p*<.0001, R^2*^=.98), such that higher melanopic irradiance was associated with higher PLR for each cognitive context (**Fig. 3**). No effect of sex, age, BMI, or time of day was detected (Executive: F_(1,11)_<1.07, *p*>.32; Emotional: F_(1,8)_<1.36, *p*>.28; Attentional: F_(1,10)_ <1.56, *p*>.24). For each task, significant differences (*p*<.002) were observed between each pair of melanopic irradiance levels.

**Figure 3.**
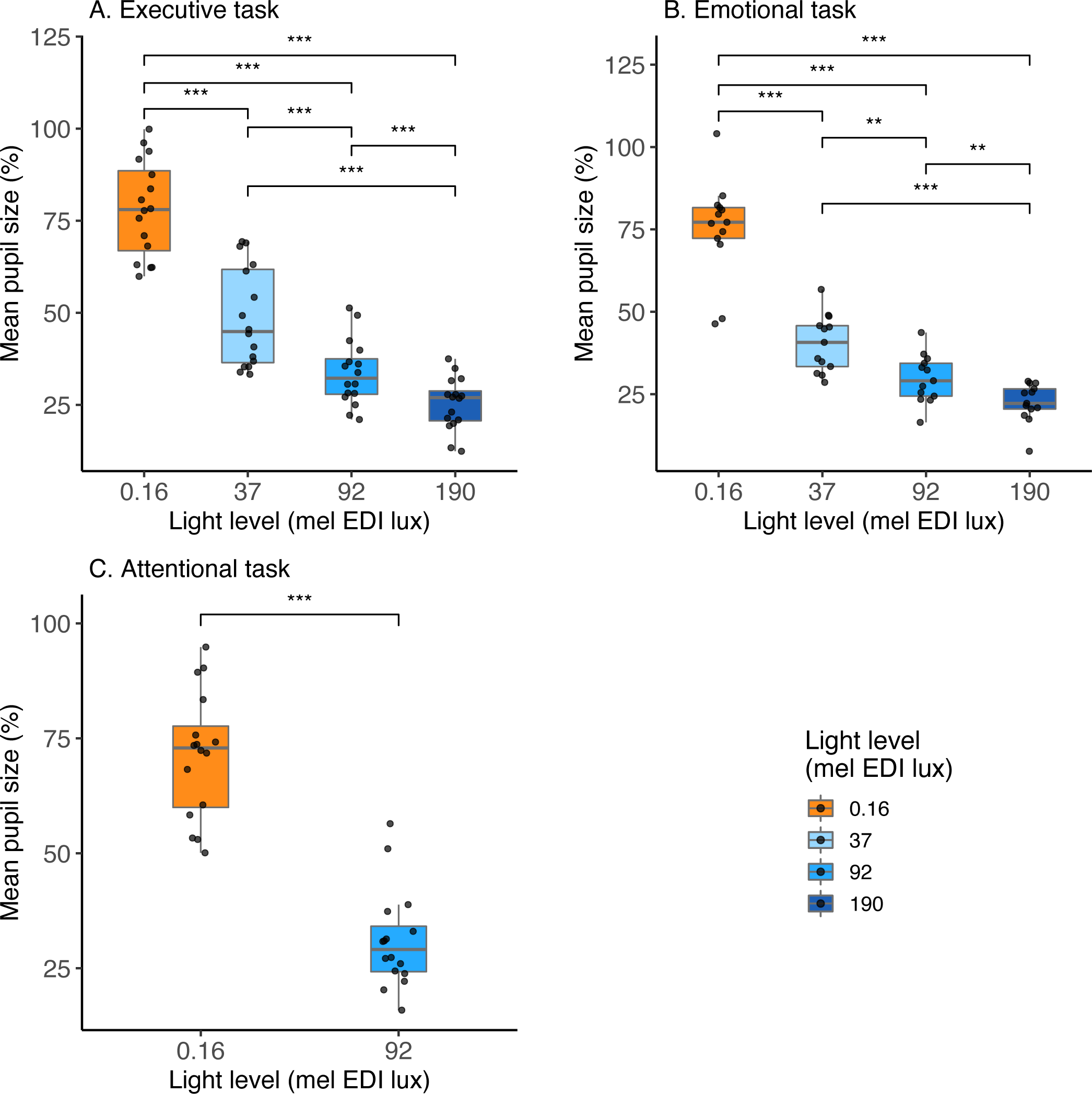
Sustained PLR across mel EDI lux levels and tasks. Darkness-normalized mean pupil size under each light level (0.16, 37, 92, 190 mel EDI lux) for the executive (A), emotional (B) and attentional (C) tasks. Mel EDI lux: melanopic equivalent daytime illuminance lux. Statistical significance on the post-hoc analysis after Tukey adjustment (*** < .0001, ** < .002).

### Pupil response stability

The effect of time in protocol and light block sequence was evaluated on the pupil response stability in each task separately (**Fig. 4**). Sustained PLR was compared during the first versus last block of each melanopic irradiance level (**Fig. 4A, 4C, 4E**). GLMM analysis revealed a significant interaction between irradiance level and light block order while controlling for sex, age, BMI, and time of day for the executive task only (F_(3,33)_=4.8, *p*=.007, R^2*^=.3). No significant interactions between melanopic irradiance level and block order were detected for the emotional and attentional tasks (F_(3,26)_=1.03, *p*=.4, and F_(1,16)_=1.96, *p*=.18, respectively). Post-hoc analyses compared light block order against each irradiance level in the executive task, but also in the other tasks. These post-hoc analyses yielded significant differences between the first and last light blocks for 0.16 and 37 mel EDI lux levels, for all three tasks [0.16 mel EDI lux: t_33_=-2.26, *p*=.031 (Executive), t_26_=-2.37, *p*=.025 (Emotional), t_16_=-3.11, *p*=.007 (Attentional); 37 mel EDI lux: t_33_=-3.26, *p*=.003 (Executive), t_26_=-2.66, *p*=.013 (Emotional)]. No significant differences were detected between the first and last blocks for the 92 and 190 mel EDI lux conditions for all three tasks (*p*>.17). Taken together, these results revealed smaller PLR over the last block compared to the first block when considering lower melanopic irradiance levels.

**Figure 4.**
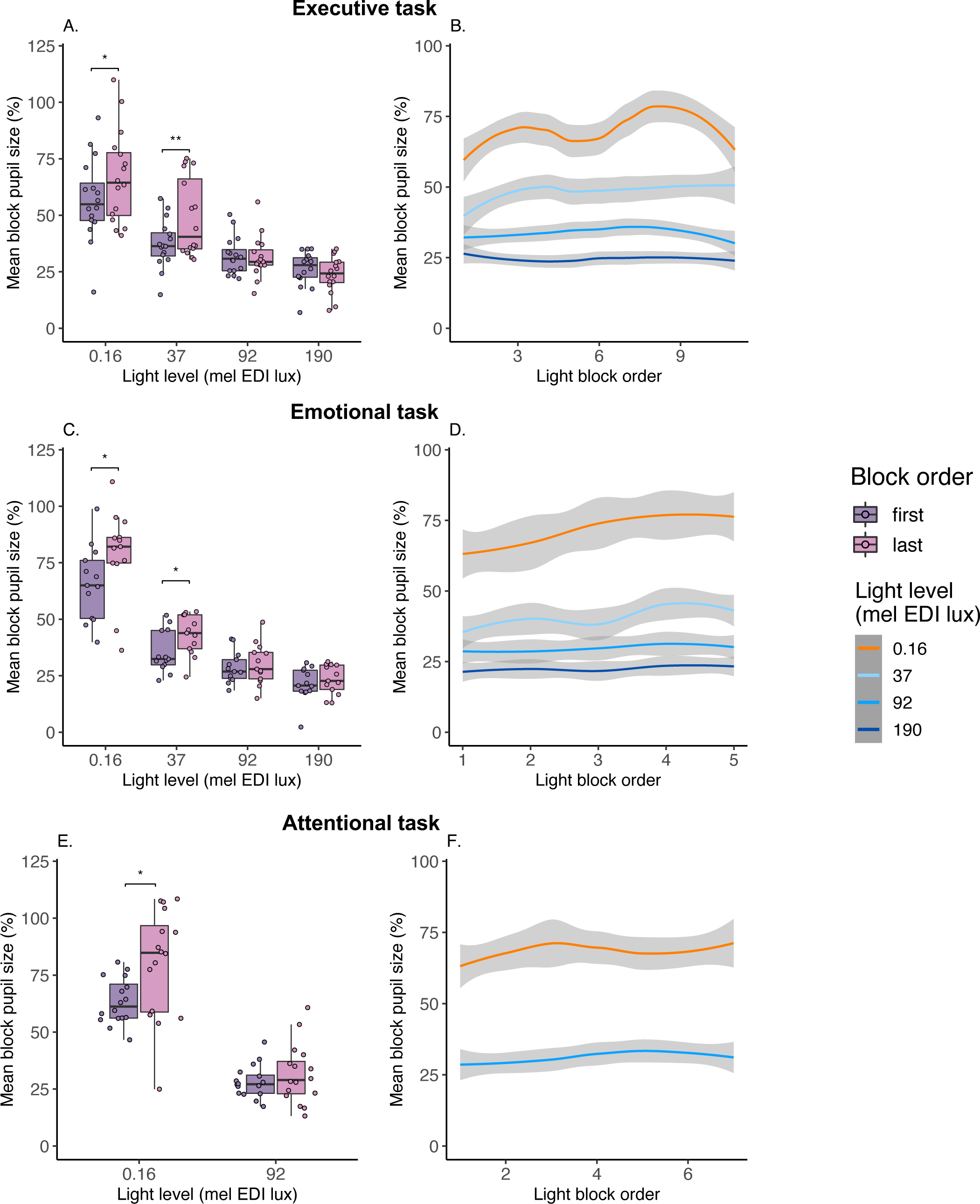
PLR stability across light blocks. **Left**: Darkness-normalized mean pupil size during the first and last block of each light level for all three cognitive tasks. Statistical significance on the post-hoc analysis after Tukey adjustment (** < .005, * < .05). **Right**: Complete evolution of darkness-normalized mean block pupil size under each light level for the three cognitive tasks. Shaded grey areas surrounding the average curves indicate the standard error. **A-B**. Executive task, 0.16, 37, 92, 190 mel EDI lux. **C-D**. Emotional task 0.16, 37, 92, 190 mel EDI lux. **E-F**. Attentional task, 0.16, 92 mel EDI lux. Mel EDI lux: melanopic equivalent daytime illuminance lux.

### Frequency analysis of sustained pupil response

To better characterize the influence of melanopic irradiance level on pupil response variability, exploratory frequency-domain analyses were performed on the PSD of sustained PLR in the 0.5-4Hz frequency range for each task separately (**Fig. 5**). The GLMM with total power as dependent variable yielded a significant main effect of melanopic irradiance level for all three tasks (Executive: F_(3,13)_=23.7, *p*<.0001, R^2*^=.85; Emotional: F_(3,16)_=130.75, *p*<.0001, R^2*^=.96; Attentional: F_(1,5)_=40.2, *p*=.0011, R^2*^=.89), such that greater PSD is observed under lower melanopic irradiance level. Post-hoc analyses on irradiance levels highlighted significant differences in the emotional and attentional tasks for all light levels (*p*<.0011). Interestingly, for the executive task, all light levels were significantly different from one another (*p*<.041), except for the lowest and the highest melanopic irradiance level ([0.16 – 37 mel EDI lux]: t_13_=2.22, *p*=.17; [92 – 190 mel EDI lux]: t_13_=2.59, *p*=.093).

**Figure 5.**
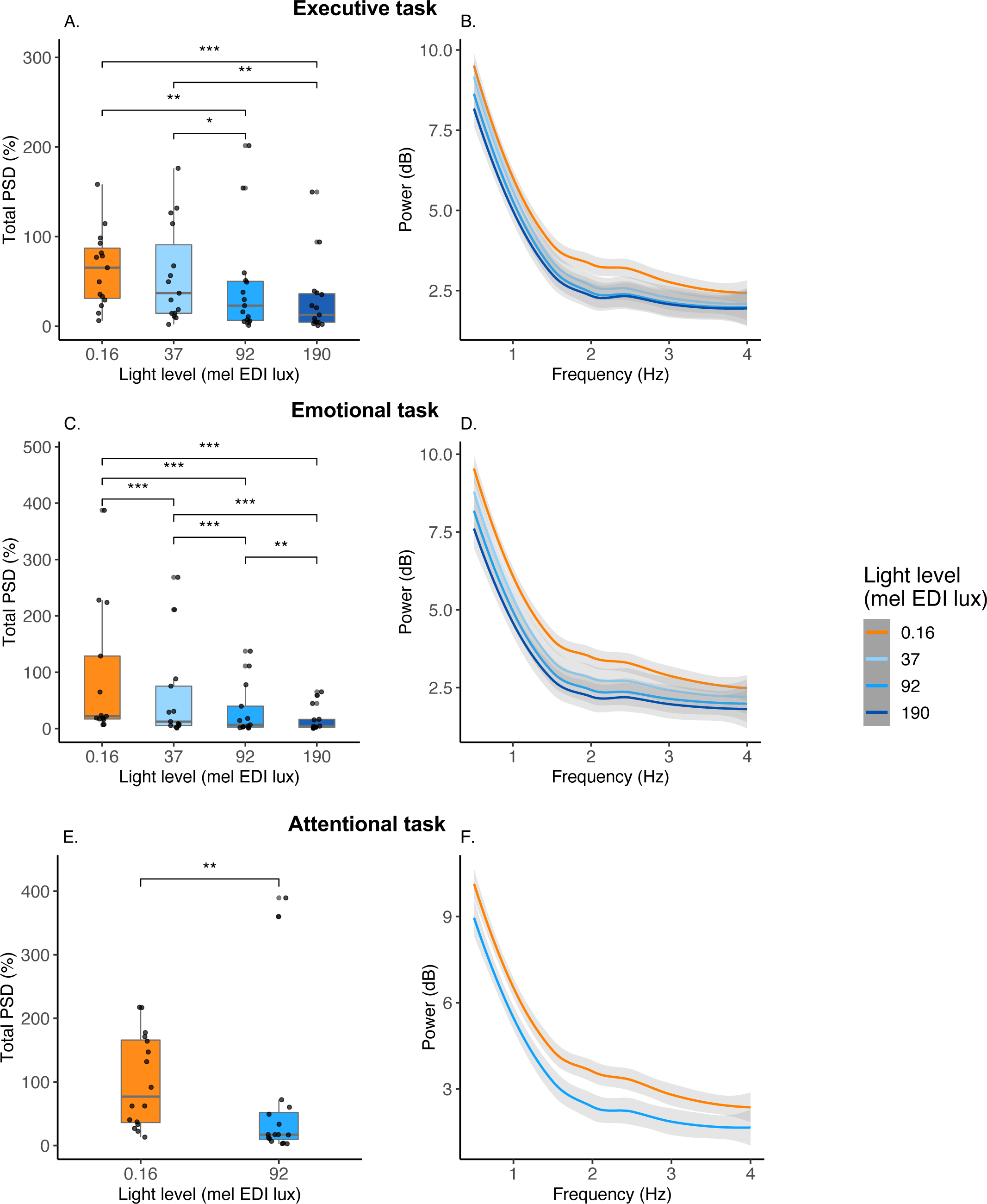
Pupil size oscillations across light conditions. **Left:** Dark-normalized total power spectrum density of pupil size across light levels for all three cognitive tasks. Statistical significance on the post-hoc analysis after Tukey adjustment (*** < .0001, ** < .005, * < .05). **Right:** Dark-normalized power spectrum density under various light conditions for all three cognitive tasks. Shaded grey areas surrounding the average curves indicate the standard error range across participants for each light condition. **A-B**. Executive task, 0.16, 37, 92, 190 mel EDI lux. **C-D**. Emotional task 0.16, 37, 92, 190 mel EDI lux. **E-F**. Attentional task, 0.16, 92 mel EDI lux. Mel EDI lux: melanopic equivalent daytime illuminance lux.

## Discussion

We characterized sustained PLR under different cognitive domain and light conditions. Thirteen to 16 healthy young individuals completed 3 different cognitive tasks while in an MRI apparatus and being exposed to repeated alternating short light blocks of different melanopic irradiance, as indexed by mel EDI lux. We replicated that a higher melanopic irradiance level leads to a smaller sustained pupil size. Our analyses further show that this effect is consistent across all three cognitive domains, taken separately. Across each task, PLR was stable in time for higher irradiance levels (92 and 190 mel EDI lux), while PLR decreased from the first until the last block for lower irradiance ones (0.16 and 37 mel EDI lux). To further characterize the variability of the sustained PLR in terms of slow oscillation power density, we found that PLR variability within the 0.5 to 4Hz range was lower under higher melanopic irradiance. Finally, PLR may vary between the executive and both the attentional and emotional tasks irrespectively of the current light condition.

Although protocols can vary substantially, PLR assessments most often used exposures lasting one to a few minutes separated by a relatively long period of darkness or constant exposure, which allows for readaptation of all retinal photoreceptors to the ambient light level [e.g. (Daneault et al., 2012; Prayag et al., 2019; Rukmini et al., 2017)]. So far, it was unclear whether a neuroimaging protocol where light exposures are shorter (1 minute or less) and interleaved with short 10 to 20s periods in darkness would affect PLR. We used light levels similar to many previous PLR studies (Daneault et al., 2012; Prayag et al., 2019; Rukmini et al., 2017) and first confirm, as expected, that PLR is stronger under higher melanopic irradiance.

Light adaptation mechanisms affect retinal photoreceptors and their associated neural circuits over the course of light exposure to optimize their sensitivity according to the ambient light levels (Lucas et al., 2012). Since the executive task was always administered first – following 45 minutes under a dim light – while the other two tasks were pseudo-randomly following the executive task and its light exposures, one could have expected a progressive reduction in PLR across the protocol. We find however a statistical trend suggesting a reduced PLR during the executive task compared to the following emotional and attentional tasks, while there was no difference between the latter two tasks. The dim light condition and associated adaptation preceding the executive task may have therefore contributed to a lower sensitivity to light, although we do not observe a difference in baseline pupil size under complete darkness between tasks (*p*>.27; data not shown). The nature of the ongoing cognitive task may also have influenced PLR since pupil size is influenced by the cognitive context (Joshi & Gold, 2020). The trends we observe should be verified in a larger sample allowing to fully separate light history from the cognitive context. We emphasize that, despite overall changes in PLR across tasks, we found no evidence of an existing relationship between tasks and irradiance light levels. The putative light adaptation mechanisms do not seem therefore to significantly affect the relative variations in PLR with irradiance levels across tasks. It remains therefore appropriate to compare tasks with respect to the relative changes in corneal irradiance levels, despite the fact that the attentional task only included two irradiance levels preventing a complete comparison across the tasks.

Overall, our results emphasize that, together with cognitive context, recent light history may influence PLR assessment and should therefore be carefully taken into account. Our findings question the appropriateness of the light history standardization period that was implemented in the protocol, with 5 min of bright (1000 lux) light exposure followed by 45 min under dim light (<10 lux). Both human and rodent data suggest that melanopsin-dependent photoreception is the main driver of NIF responses to light under more naturalistic conditions, i.e. not following dim light or dark adaptation (Lucas et al., 2012). Standardization using higher ambient light levels (and potentially over shorter periods of time) would simplify experimental procedures and reduce the glare effect most participants experience during the first block(s) of exposure to light while maintaining, or potentially improving, the sensitivity to melanopsin-driven photoreception. This warrants future investigations comparing different pre-recording standardization procedures.

The fact that sustained PLR was reduced from the first to the last block of light for mel EDI lux levels inferior and equal to 37 mel EDI lux suggests that light adaptation did affect photoreceptors sensitivity over time within a task. Since rods and cones contribute more than melanopsin-dependent photoreception to PLR at lower light levels (Do et al., 2009; Gooley et al., 2012; Lucas et al., 2003; McDougal & Gamlin, 2010), we suspect that light adaptation of either rods or cones, or both, contributed to a reduction in PLR over time. At higher melanopic irradiance levels, PLR is more heavily driven by the intrinsic melanopsin-dependent photoreception of ipRGCs which show a much slower adaptation to the ambient light level (Gooley et al., 2012). Our data support therefore that the intrinsic photoreception of ipRGCs drives the relatively stable PLR we observe from the beginning until the end of each task for melanopic irradiance of ∼90 mel EDI lux or higher.

Studies assessing neural correlates of NIF effects light often used repeated alternating short light exposures with varying irradiances and spectral compositions (Daneault et al., 2014; Gaggioni et al., 2014; McGlashan et al., 2021; Vandewalle et al., 2011; Vandewalle et al., 2010). Although light characteristic descriptions were not always exhaustive, these studies seem to have mostly used melanopic irradiance levels higher than ∼90 and up to ∼330 mel EDI lux, except for a few cases which also included a light condition of ∼20 mel EDI lux in addition to higher irradiances (Daneault et al., 2014; Vandewalle et al., 2011; Vandewalle et al., 2010). Hence, the present results support that these studies did not suffer from important bias related to photoreceptors adaptation over short exposures (1 min or less) separated by brief periods of darkness. The fMRI data associated with the present study will nevertheless need to account for potential photoreceptor adaptation, if not in their analysis, at least in the interpretation of the results.

The finding that higher melanopic irradiance is associated with lower power density over the 0.5-4Hz frequency band may appear surprising as it shows that pupil response oscillations (over 0.25 to 2 seconds periods) were less important at higher irradiances. This could indeed imply a reduced tonic activity of the LC when light is known to stimulate alertness and higher alertness is associated with higher LC activity (Aston-Jones & Bloom, 1981). Previous research reported, however, higher power densities of fast pupil size oscillation under lower background luminance (Nakayama & Shimizu, 2021; Nguyen et al., 2022; Peysakhovich et al., 2015). In addition, increasing arousal is associated first with a higher rate of tonic LC firing, and then with a switch to a phasic firing of the LC activity (Aston-Jones & Bloom, 1981). The reduced power density could therefore be the consequence of a change in the firing pattern of the LC. It could also result from a change in the frequency of the fast oscillations in pupil diameter outside the frequency band we considered. In a companion paper, we focus on transient changes in pupil size that are arguably related to the phasic activity of the LC [Campbell et al. bioRxiv]. Yet, the transient dilations of the pupil are induced by the auditory stimulations included in the cognitive task recorded in fMRI (Murphy et al., 2014). We are therefore not in a position to assess a putative LC phasic activity that would not be related to the sensory stimulations and contribute to changes in power density of pupil size variations. Importantly also, since pupil size is not governed by the LC but rather influenced by it, our findings could be driven by other brain structures.

We stress that our research bears some limitations. First, we included 4 distinct melanopic irradiance levels, preventing the establishment of a true action spectrum of the PLR under the conditions of our experiments (Mure, 2021). In addition, two distinct spectral qualities or colors were used so that visual responses to the perception of a control orange exposure could be subtracted from the response to the active blue-enriched polychromatic light in the analyses of fMRI data (Daneault et al., 2014; Vandewalle et al., 2007). This implies that part of our findings regarding PLR may be related to spectral differences and not only to irradiance levels. Given the relative homogeneity of our findings across tasks, we remain confident that the significant differences we find are robust. Finally, as our primary interest was to relate sustained PLR to melanopic irradiance levels, we did not consider the initial phasic portion of PLR over the first 2 seconds of the exposure which is known to rely more heavily on rods and/or cone photoreception (Gooley et al., 2012). Hence, we cannot exclude that changes in the sensitivity of these photoreceptors impacted this initial portion of the PLR.

Light is an important environmental factor affecting brain functions, behavior, health and well-being. Given the expansion of artificial light usage, a detailed understanding of its NIF impacts is timely. With this study, we emphasize that PLR is an easy readout of one of the multiple NIF effects of light and that it can be used as a window to the underlying brain mechanisms. We provided information on the association between sustained PLR and melanopic irradiance levels under the specific context of an fMRI protocol. We show that depending on the experimental conditions, photoreceptors adaptation may or may not significantly affect the NIF responses of interest. We further suggest that the light adaptation period may influence PLR. Since PLR can be easily characterized across different species, our results will contribute to the translation of animal findings to human beings and vice-versa (Lucas et al., 2003).

## Conflict of interest

The authors declare no conflict of interests.

## Author contributorship

E.B., I.C., G.V. designed the research.

E.B., I.C., R.S., I.P. and J.F.B.A. acquired the data.

A.B., E.K., N.M., P.T., S.S. provided valuable insights while acquiring, interpreting, and discussing the data.

E.B. analysed the data supervised by G.V. E.B., H.J., G.V. wrote the paper.

All authors edited and approved the final version of the manuscript.

## Data and code availability statement

The processed data and analysis scripts supporting the results included in this manuscript are publicly available via the following open repository: https://gitlab.uliege.be/CyclotronResearchCentre/Public/xxxx (the repository will be created following acceptance of the paper / upon request of the editor)

## Acknowledgments

We thank A. Claes, C. Degueldre, C. Hagelstein, G. Hammad, B. Herbillon, P. Hawotte, E. Lambot, B. Lauricella, A. Luxen and E. Salmon for their help in the different steps of the project.

## Fundings

This project has received funding from the European Union’s Horizon 2020 research and innovation programme under the Marie Skłodowska-Curie grant agreement No 860613 (LIGHTCAP project). This study was also supported by the Belgian Fonds de la Recherche Scientifique (FRS-FNRS; CDR J.0222.20), the Fondation Léon Frédéricq, ULiège - U. Maastricht Imaging Valley, ULiège-Valeo Innovation Chair “Health and Well-Being in Transport” and Safran (LIGHT-CABIN project), the European Regional Development Fund (Biomed-hub), and Siemens Healthineers. None of these funding sources had any impact on the design of the study nor on the interpretation of the findings. E.B. is supported by ULiège - U. Maastricht Imaging Valley. I.C., N.M., E.K. C.P., and G.V. are/were supported by the FNRS. R.S and F.B are supported by the European Union’s Horizon 2020 research and innovation programme under the Marie Skłodowska-Curie grant agreement No 860613 (LIGHTCAP project). I.P. is supported by the FNRS and the GIGA Doctoral School for Health Sciences of ULiège. AB is supported by Synergia Medial SA and the Walloon Region (Industrial Doctorate Program, convention n°8193). S.S. was supported by ULiège-Valeo Innovation Chair “Health and Well-Being in Transport” and Safran and by Siemens Healthineers. N.M. and E.K. were supported in part by the Fondation Recherche Alzheimer (SOA-FRA).

## Notes

### Competing Interest Statement

The authors have declared no competing interest.

### Summary of Updates

This version of the manuscript includes the mention and reference to the companion paper of Campbell et al. (bioRxiv).

## References

Aston-Jones, G., & Bloom, F. E. (1981, Aug). Norepinephrine-containing locus coeruleus neurons in behaving rats exhibit pronounced responses to non-noxious environmental stimuli. J Neurosci, 1(8), 887–900. https://doi.org/10.1523/jneurosci.01-08-00887.1981

Banse, R., & Scherer, K. R. (1996, Mar). Acoustic profiles in vocal emotion expression. J Pers Soc Psychol, 70(3), 614–636. https://doi.org/10.1037//0022-3514.70.3.614

Beck, A. T., Epstein, N., Brown, G., & Steer, R. A. (1988, Dec). An inventory for measuring clinical anxiety: psychometric properties. J Consult Clin Psychol, 56(6), 893–897. https://doi.org/10.1037//0022-006x.56.6.893

Beck, A. T., Ward, C. H., Mendelson, M., Mock, J., & Erbaugh, J. (1961, Jun). An inventory for measuring depression. Arch Gen Psychiatry, 4, 561–571. https://doi.org/10.1001/archpsyc.1961.01710120031004

Berson, D. M., Dunn, F. A., & Takao, M. (2002, Feb 8). Phototransduction by retinal ganglion cells that set the circadian clock. Science, 295(5557), 1070–1073. https://doi.org/10.1126/science.1067262

Brainard, G. C., Hanifin, J. P., Greeson, J. M., Byrne, B., Glickman, G., Gerner, E., & Rollag, M. D. (2001, Aug 15). Action spectrum for melatonin regulation in humans: evidence for a novel circadian photoreceptor. J Neurosci, 21(16), 6405–6412. https://doi.org/10.1523/jneurosci.21-16-06405.2001

Buysse, D. J., Reynolds, C. F., 3rd, Monk, T. H., Berman, S. R., & Kupfer, D. J. (1989, May). The Pittsburgh Sleep Quality Index: a new instrument for psychiatric practice and research. Psychiatry Res, 28(2), 193-213. https://doi.org/10.1016/0165-1781(89)90047-4

Cajochen, C., Münch, M., Kobialka, S., Kräuchi, K., Steiner, R., Oelhafen, P., Orgül, S., & Wirz-Justice, A. (2005, Mar). High sensitivity of human melatonin, alertness, thermoregulation, and heart rate to short wavelength light. J Clin Endocrinol Metab, 90(3), 1311–1316. https://doi.org/10.1210/jc.2004-0957

Collette, F., Hogge, M., Salmon, E., & Van der Linden, M. (2005, Apr 28). Exploration of the neural substrates of executive functioning by functional neuroimaging. Neuroscience, 139(1), 209–221. https://doi.org/10.1016/j.neuroscience.2005.05.035

Daneault, V., Hébert, M., Albouy, G., Doyon, J., Dumont, M., Carrier, J., & Vandewalle, G. (2014, Jan 1). Aging reduces the stimulating effect of blue light on cognitive brain functions. Sleep, 37(1), 85–96. https://doi.org/10.5665/sleep.3314

Daneault, V., Vandewalle, G., Hébert, M., Teikari, P., Mure, L. S., Doyon, J., Gronfier, C., Cooper, H. M., Dumont, M., & Carrier, J. (2012, Jun). Does pupil constriction under blue and green monochromatic light exposure change with age? J Biol Rhythms, 27(3), 257–264. https://doi.org/10.1177/0748730412441172

Do, M. T., Kang, S. H., Xue, T., Zhong, H., Liao, H. W., Bergles, D. E., & Yau, K. W. (2009, Jan 15). Photon capture and signalling by melanopsin retinal ganglion cells. Nature, 457(7227), 281–287. https://doi.org/10.1038/nature07682

Fisk, A. S., Tam, S. K. E., Brown, L. A., Vyazovskiy, V. V., Bannerman, D. M., & Peirson, S. N. (2018). Light and Cognition: Roles for Circadian Rhythms, Sleep, and Arousal. Front Neurol, 9, 56. https://doi.org/10.3389/fneur.2018.00056

Gaggioni, G., Maquet, P., Schmidt, C., Dijk, D. J., & Vandewalle, G. (2014). Neuroimaging, cognition, light and circadian rhythms. Front Syst Neurosci, 8, 126. https://doi.org/10.3389/fnsys.2014.00126

Gamlin, P. D., McDougal, D. H., Pokorny, J., Smith, V. C., Yau, K. W., & Dacey, D. M. (2007, Mar). Human and macaque pupil responses driven by melanopsin-containing retinal ganglion cells. Vision Res, 47(7), 946–954. https://doi.org/10.1016/j.visres.2006.12.015

Gooley, J. J., Ho Mien, I., St Hilaire, M. A., Yeo, S. C., Chua, E. C., van Reen, E., Hanley, C. J., Hull, J. T., Czeisler, C. A., & Lockley, S. W. (2012, Oct 10). Melanopsin and rod-cone photoreceptors play different roles in mediating pupillary light responses during exposure to continuous light in humans. J Neurosci, 32(41), 14242–14253. https://doi.org/10.1523/jneurosci.1321-12.2012

Güler, A. D., Ecker, J. L., Lall, G. S., Haq, S., Altimus, C. M., Liao, H. W., Barnard, A. R., Cahill, H., Badea, T. C., Zhao, H., Hankins, M. W., Berson, D. M., Lucas, R. J., Yau, K. W., & Hattar, S. (2008, May 1). Melanopsin cells are the principal conduits for rod-cone input to non-image-forming vision. Nature, 453(7191), 102–105. https://doi.org/10.1038/nature06829

Hattar, S., Kumar, M., Park, A., Tong, P., Tung, J., Yau, K. W., & Berson, D. M. (2006, Jul 20). Central projections of melanopsin-expressing retinal ganglion cells in the mouse. J Comp Neurol, 497(3), 326–349. https://doi.org/10.1002/cne.20970

Horne, J. A., & Ostberg, O. (1976). A self-assessment questionnaire to determine morningness-eveningness in human circadian rhythms. Int J Chronobiol, 4(2), 97–110.

Jaeger, B. C., Edwards, L. J., Das, K., & Sen, P. K. (2017, 2017/04/26). An R2 statistic for fixed effects in the generalized linear mixed model. Journal of Applied Statistics, 44(6), 1086-1105. https://doi.org/10.1080/02664763.2016.1193725

Johns, M. W. (1991, Dec). A new method for measuring daytime sleepiness: the Epworth sleepiness scale. Sleep, 14(6), 540–545. https://doi.org/10.1093/sleep/14.6.540

Joshi, S., & Gold, J. I. (2020). Pupil Size as a Window on Neural Substrates of Cognition. Trends in Cognitive Sciences, 24(6), 466–480. https://doi.org/10.1016/j.tics.2020.03.005

Joshi, S., Li, Y., Kalwani, R. M., & Gold, J. I. (2016, Jan 6). Relationships between Pupil Diameter and Neuronal Activity in the Locus Coeruleus, Colliculi, and Cingulate Cortex. Neuron, 89(1), 221–234. https://doi.org/10.1016/j.neuron.2015.11.028

Lok, R., Smolders, K., Beersma, D. G. M., & de Kort, Y. A. W. (2018, Dec). Light, Alertness, and Alerting Effects of White Light: A Literature Overview. J Biol Rhythms, 33(6), 589–601. https://doi.org/10.1177/0748730418796443

Lucas, R. J., Hattar, S., Takao, M., Berson, D. M., Foster, R. G., & Yau, K. W. (2003, Jan 10). Diminished pupillary light reflex at high irradiances in melanopsin-knockout mice. Science, 299(5604), 245–247. https://doi.org/10.1126/science.1077293

Lucas, R. J., Lall, G. S., Allen, A. E., & Brown, T. M. (2012). Chapter 1 - How rod, cone, and melanopsin photoreceptors come together to enlighten the mammalian circadian clock. In A. Kalsbeek, M. Merrow, T. Roenneberg, & R. G. Foster (Eds.), Progress in Brain Research (Vol. 199, pp. 1-18). Elsevier. https://doi.org/10.1016/B978-0-444-59427-3.00001-0

Lucas, R. J., Peirson, S. N., Berson, D. M., Brown, T. M., Cooper, H. M., Czeisler, C. A., Figueiro, M. G., Gamlin, P. D., Lockley, S. W., O’Hagan, J. B., Price, L. L. A., Provencio, I., Skene, D. J., & Brainard, G. C. (2014, 2014/01/01/). Measuring and using light in the melanopsin age. Trends in Neurosciences, 37(1), 1–9. https://doi.org/10.1016/j.tins.2013.10.004

Mathôt, S., Schreij, D., & Theeuwes, J. (2012, 2012/06/01). OpenSesame: An open-source, graphical experiment builder for the social sciences. Behavior Research Methods, 44(2), 314–324. https://doi.org/10.3758/s13428-011-0168-7

McDougal, D. H., & Gamlin, P. D. (2010, Jan). The influence of intrinsically-photosensitive retinal ganglion cells on the spectral sensitivity and response dynamics of the human pupillary light reflex. Vision Res, 50(1), 72–87. https://doi.org/10.1016/j.visres.2009.10.012

McGlashan, E. M., Poudel, G. R., Jamadar, S. D., Phillips, A. J. K., & Cain, S. W. (2021). Afraid of the dark: Light acutely suppresses activity in the human amygdala. PLoS One, 16(6), e0252350. https://doi.org/10.1371/journal.pone.0252350

Megemont, M., McBurney-Lin, J., & Yang, H. (2022, 2022/02/02). Pupil diameter is not an accurate real-time readout of locus coeruleus activity. eLife, 11, e70510. https://doi.org/10.7554/eLife.70510

Mure, L. S. (2021). Intrinsically Photosensitive Retinal Ganglion Cells of the Human Retina. Front Neurol, 12, 636330. https://doi.org/10.3389/fneur.2021.636330

Murphy, P. R., O’Connell, R. G., O’Sullivan, M., Robertson, I. H., & Balsters, J. H. (2014, Aug). Pupil diameter covaries with BOLD activity in human locus coeruleus. Hum Brain Mapp, 35(8), 4140-4154. https://doi.org/10.1002/hbm.22466

Nakayama, M., & Shimizu, Y. (2021). Frequency Analysis of Task Evoked Pupillary Response and Eye Movement. In M. Nakayama & Y. Shimizu (Eds.), Pupil Reactions in Response to Human Mental Activity (pp. 89-103). Springer Singapore. https://doi.org/10.1007/978-981-16-1722-5_7

Nguyen, K. T., Liang, W. K., Juan, C. H., & Wang, C. A. (2022, Jun). Time-frequency analysis of pupil size modulated by global luminance, arousal, and saccade preparation signals using Hilbert-Huang transform. Int J Psychophysiol, 176, 89–99. https://doi.org/10.1016/j.ijpsycho.2022.03.011

Peysakhovich, V., Causse, M., Scannella, S., & Dehais, F. (2015, Jul). Frequency analysis of a task-evoked pupillary response: Luminance-independent measure of mental effort. Int J Psychophysiol, 97(1), 30–37. https://doi.org/10.1016/j.ijpsycho.2015.04.019

Prayag, A., Jost-Boissard, S., Avouac, P., Dumortier, D., & Gronfier, C. (2019, 03/05). Dynamics of Non-visual Responses in Humans: As Fast as Lightning? Frontiers in Neuroscience, 13. https://doi.org/10.3389/fnins.2019.00126

Rosenthal, N. (1984). Seasonal pattern assessment questionnaire. Journal of Affective Disorders.

Rukmini, A. V., Milea, D., Aung, T., & Gooley, J. J. (2017, Mar 7). Pupillary responses to short-wavelength light are preserved in aging. Sci Rep, 7, 43832. https://doi.org/10.1038/srep43832

Stevens, A. A., Skudlarski, P., Gatenby, J. C., & Gore, J. C. (2000, Jun). Event-related fMRI of auditory and visual oddball tasks. Magn Reson Imaging, 18(5), 495–502. https://doi.org/10.1016/s0730-725x(00)00128-4

Thapan, K., Arendt, J., & Skene, D. J. (2001, Aug 15). An action spectrum for melatonin suppression: evidence for a novel non-rod, non-cone photoreceptor system in humans. J Physiol, 535(Pt 1), 261–267. https://doi.org/10.1111/j.1469-7793.2001.t01-1-00261.x

Vandewalle, G., Archer, S. N., Wuillaume, C., Balteau, E., Degueldre, C., Luxen, A., Dijk, D. J., & Maquet, P. (2011, Jun). Effects of light on cognitive brain responses depend on circadian phase and sleep homeostasis. J Biol Rhythms, 26(3), 249–259. https://doi.org/10.1177/0748730411401736

Vandewalle, G., Maquet, P., & Dijk, D. J. (2009, Oct). Light as a modulator of cognitive brain function. Trends Cogn Sci, 13(10), 429–438. https://doi.org/10.1016/j.tics.2009.07.004

Vandewalle, G., Schmidt, C., Albouy, G., Sterpenich, V., Darsaud, A., Rauchs, G., Berken, P. Y., Balteau, E., Degueldre, C., Luxen, A., Maquet, P., & Dijk, D. J. (2007, Nov 28). Brain responses to violet, blue, and green monochromatic light exposures in humans: prominent role of blue light and the brainstem. PLoS One, 2(11), e1247. https://doi.org/10.1371/journal.pone.0001247

Vandewalle, G., Schwartz, S., Grandjean, D., Wuillaume, C., Balteau, E., Degueldre, C., Schabus, M., Phillips, C., Luxen, A., Dijk, D. J., & Maquet, P. (2010, Nov 9). Spectral quality of light modulates emotional brain responses in humans. Proc Natl Acad Sci U S A, 107(45), 19549–19554. https://doi.org/10.1073/pnas.1010180107

